# Acoustic light-sheet microscopy

**DOI:** 10.1101/2021.08.20.457051

**Authors:** S. Wunderl, A. Ishijima, E. A. Susaki, Z. Xu, H. Song, H. Zha, T. Azuma, I. Sakuma, H. Fukuoka, E. Okada, H. R. Ueda, S. Takagi, K. Nakagawa

## Abstract

Light-sheet imaging of 3D objects with high spatial resolution remains an open challenge because of the trade-off between field-of-view (FOV) and axial resolution originating from the diffraction of light. We developed acoustic light-sheet microscopy (acoustic LSM), which actively manipulates the light propagation inside a large sample to obtain wide-field microscopic images deep inside a target. By accurately coupling a light-sheet illumination pulse into a planar acoustic pulse, the light-sheet can be continuously guided over large distances. We imaged a fluorescence-labeled transparent mouse brain for the FOVs of 19.3 × 12.4 mm^2^ and 9.7 × 5.9 mm^2^ with resolved microstructures and single cells deep inside the brain. Acoustic LSM creates new opportunities for the application of light-sheet in the field of industry to basic science.

**One Sentence Summary:** An acoustic-optical method overcomes a trade-off between field-of-view and axial resolution in light-sheet microscopy.

## Main Text

Since the discovery of the cell as the basic unit of living organisms, people have been seeking a way to map the whole biological system with single-cell resolution (*1*–*4*). Light-sheet fluorescence microscopy (LSFM) with tissue clearing methods is an indispensable tool for this purpose (*5*–*10*). The basic principle of LSFM is to optically section a 3D object with a thin light-sheet (*11*–*13*). However, there is a trade-off between imaging field-of-view (FOV) and axial resolution in conventional LSFM systems originating from the diffraction of the light, where the achievable FOV is constrained by the thickness of the light-sheet. A large field of view in imaging requires the sacrifice of the axial resolution, which additionally results in the degradation of lateral resolution due to the accumulation of the spatial profile of samples in the axial direction. This fundamental limitation is an open challenge to achieve wide-field single-cell resolved imaging of large organs. Although multi-view imaging (*18*, *19*), bessel beams (*14*–*16*), and axial-scanning systems (*20, 21*) can enlarge the FOV while maintaining a high axial resolution, compromises like repetitive measurements or complicated mechanical stitching procedures must still be made. This can cause the degradation of the sample and the mismatch in stitched images. Regardless of these recent progressions on the development of optics or devices for passively manipulating the light propagation, the inevitable compromise between length and thickness of the light-sheet remains.

In this report, we present acoustic light-sheet microscopy (acoustic LSM) to address this challenge and demonstrate its utility through wide-field fluorescence imaging of a transparent mouse brain. We propose active light manipulation for light-sheet imaging: suppress the light spreading by acoustic pressure fields inside a sample for overcoming the fundamental trade-off between FOV and axial resolution in conventional LSFM. Our technique, based on the concept of a light-guiding method with acoustic waves (*22, 23*), uses an ultrawide planar acoustic pulse to non-destructively create a large ‘acoustic-optical sheet’ by modulating the spatial profile of the refractive index in the sample (Fig. 1A). The acoustic light-sheet technique allows a thin illumination in the whole cleared organ to acquire a wide-field single-cell resolved slice image in a single shot.

**Fig. 1.**
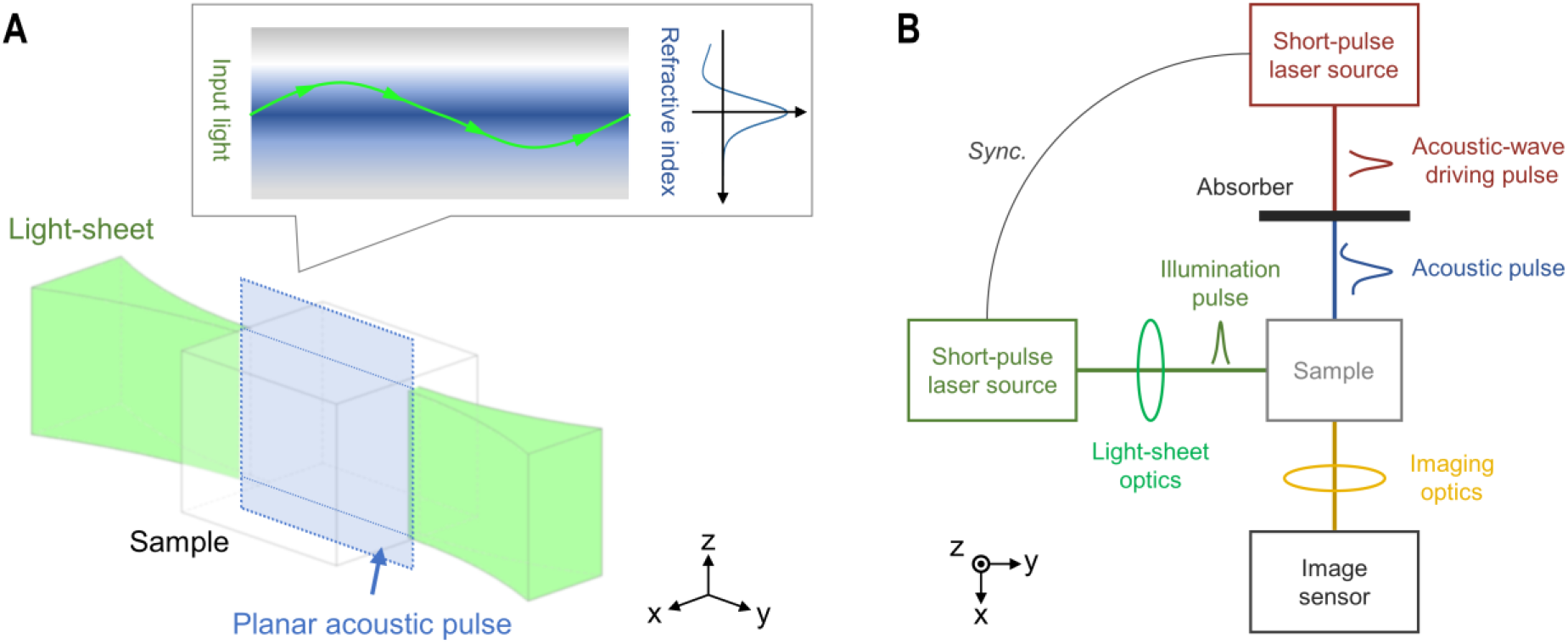
Schematic diagram of acoustic light-sheet imaging. (**A)** The principle of active light manipulation in a sample. The pressure field of a planar acoustic pulse produces a local refractive index change inside the sample. This acoustic field-based refractive index contrast creates an acoustic-optical sheet that works as optical waveguides to confine the light-sheet along the acoustic plane. (**B**) Basic configuration for acoustic light-sheet imaging. Two short-pulse lasers are used. One is an acoustic driving pulse for producing the planar acoustic pulse with a high-pressure gradient by instantaneous photon-to-phonon conversion at an absorber. The other is an illumination pulse shaped by the light-sheet optics for imaging. Since these two pulses are synchronized, and the acoustic-wave generation and propagation are reproducible events, the illumination pulse can be coupled with the acoustic pulse from the side with high accuracy. The image sensor detects the photons emitted from the sample through the imaging optics.

For realizing the proposed wide-field light-sheet microscopy, we developed an acoustic-optical system utilizing laser technologies (Fig. 1B) to meet three requirements: (a) the generation of a single acoustic pulse with a wide wavefront to achieve a wide FOV in imaging, (b) the formation of high refractive index contrast to confine light in the medium, (c) precise control of the timing between a single acoustic pulse and illumination pulse to avoid blur in the axial direction, all of which are difficult to achieve with conventional ultrasonic generators based on the vibration of piezoelectric transducers. In our method, a short laser pulse is spatially shaped to obtain the required size for the planar acoustic pulse to guide the light-sheet. The acoustic pulse is generated via light absorption at an absorber and rapid expansion of the medium. It propagates passes into the sample, where it creates a pressure field that modulates the refractive index profile. An illumination pulse synchronized with the acoustic-wave driving pulse is coupled to the planar acoustic pulse (y-z plane) to form the thin light-sheet. The emission from the sample is detected by an image sensor as in traditional light-sheet imaging. This single-shot image acquisition can be repeated with the lasers’ repetition frequency or camera’s frame rate to obtain 3D data by sliding the sample in an x-axis direction. The acoustic pulses used in our experiments resemble smoothed Friedlander curves (*24*), displaying a sharp rise of pressure (i.e. high refractive index contrast) at the leading wavefront with a following elongated negative-pressure area (fig. S1). The positive peak-pressures range up to several hundred kilopascals (depending on acoustic driving pulse laser intensity) and are low enough to avoid tissue damage (*25*).

To begin the proof-of-concept of acoustic LSM, we carried out numerical simulations of the light propagation inside acoustic pressure fields, and experimental visualizations of the light path manipulated by a planar acoustic pulse. Light propagation is simulated by the Fast Fourier transform beam propagation method (*26*, *27*). The acoustic waveguides formed in a sample were characterized by the refractive index contrast inside the medium which was estimated from experimentally measured acoustic pressure profiles (the peak pressure of 200 kPa and the positive pressure FWHM of 52.5 μm. See fig. S1–S4 and Movie S1.). When 532 nm light with a beam waist radius of 20 μm was introduced, the light travels through the sample while diverging due to the diffraction (Fig. 2A). By contrast, the light coupled to the acoustic waveguide could propagate over a long distance (35 mm shown in Fig. 2B, but not limited to). It should be noted that a large pressure gradient of laser-induced acoustic pulse enables light-guiding with a low peak-pressure acoustic field (fig. S5). In the experimental evaluation of the acoustic LSM’s light-guiding performance, we imaged the fluorescence emitted from the light-path of a light-sheet inside polyacrylamide (PAA) gel containing Rhodamine B (fig. S6 and supplementary text). Planar acoustic pulses were driven by irradiating elliptically shaped laser pulses (~ 40 × 5 mm^2^) to a water-diluted carbon colloid solution (fig. S7 and supplementary text). A 532 nm laser pulse with a pulse duration of 3-5 ns was cylindrically focused into the PAA gel sample to create a light-sheet (Rayleigh-range < 1 mm) for the excitation of fluorescence molecules. The light spread out after forming the light-sheet inside the sample (Fig. 2C) whereas the acoustic LSM technique confined the light-sheet along the acoustic field (Fig. 2D, fig. S8 and Movie S2). This effect depends on the size of a planar acoustic pulse and was confirmed up to a distance of 30 mm from the focusing point of the light-sheet (Fig. 2E).

**Fig. 2.**
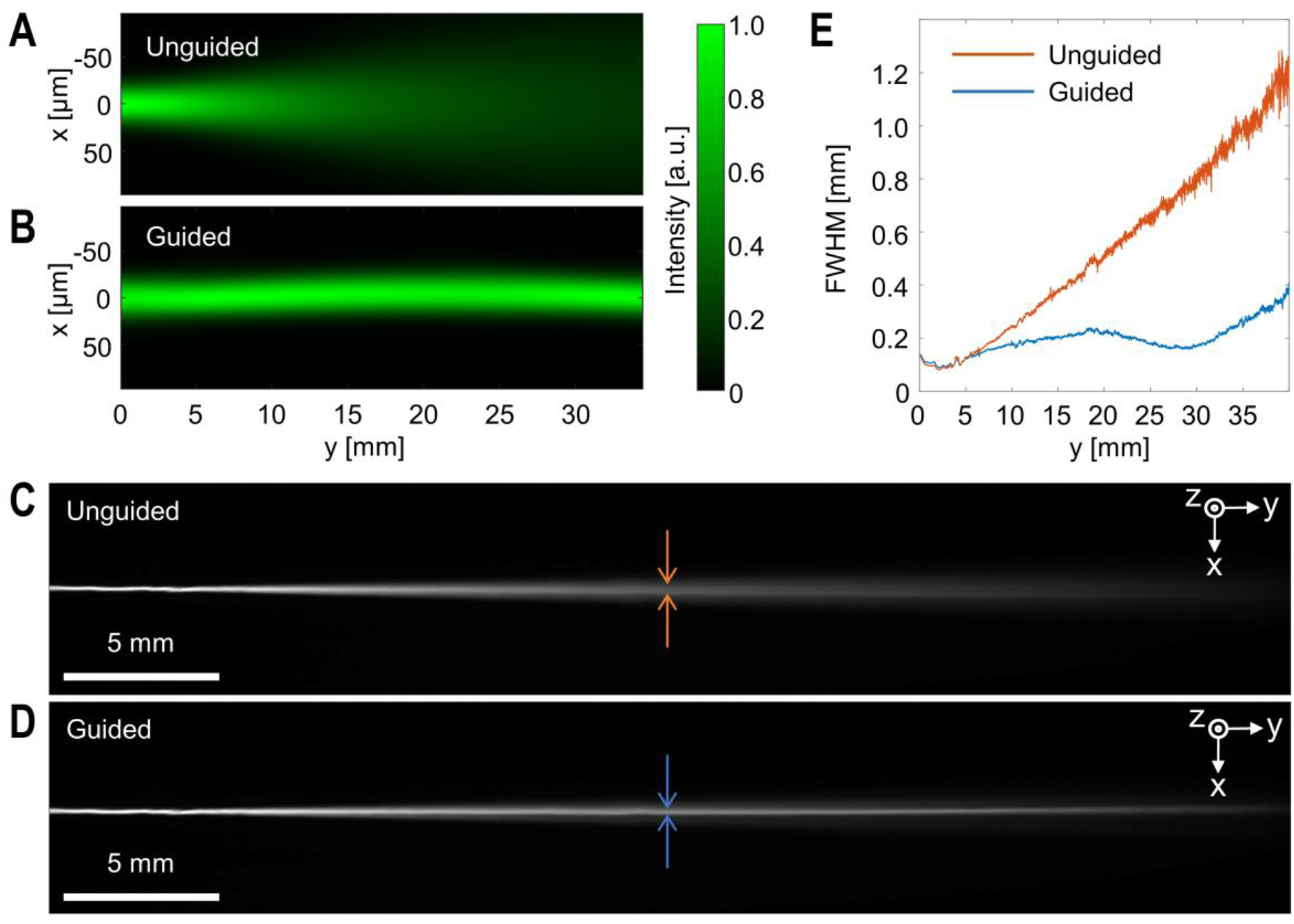
Basic performance of acoustic LSFM. **(A** and **B)** Simulation of a gaussian beam (initial beam waist radius is 20 μm, propagating from left to right) in a PAA gel for unguided and guided light propagation, respectively. The refractive index distribution produced by an acoustic pulse was set based on the pressure profile obtained in the experiments (for details, see Supplementary notes 1-4). Note that the scales of the x-axis and y-axis are micrometer and millimeter, respectively. **(C** and **D)** Fluorescence images of a Rhodamine B contained PAA gel sample captured from the z-direction. The illumination light-sheet pulses were focused at y = 2.5 mm from the left-side of the images. With the acoustic guiding technique (D), the light was confined along the acoustic-optical sheet (y-z plane). **(E)** Transitions in the FWHM of the fluorescence intensity profiles (C) and (D) in the x-axis direction.

As a demonstration of acoustic LSM, we acquired fluorescence images of a transparent mouse brain with a wide imaging FOV (Fig. 3A and fig. S9). Advancement in tissue clearing methods allows us to optically access the detailed structure and functions of large organs, e.g. primate brain (*28*). Combining the clearing methods with expansion microscopy (*29*), which physically magnifies the sample, enables super-resolution imaging of the specimens. Consequently, there is an increasing demand for an optical imaging technique to acquire whole images of large organs. In this study, we used a CUBIC-based transparent brain which cell nuclei were stained with propidium iodide (supplementary materials, materials and methods. Note that although we used the CUBIC technique here, our acoustic LSM can be applied to transparent samples prepared using other tissue clearing methods with the modification of the system based on their acoustic properties of the transparent samples.). The thickness of light-sheet and the mouse brain’s anatomical structures were captured with the FOV of 19.3 × 12.4 mm^2^ in single-shot acquisition. The classical gaussian beam properties can be observed in the unguided light-sheet image captured from the top (Fig. 3B); it showed a thin illumination area close to the Rayleigh-range of the beam waist, and it spread when advancing further. In contrast, the guided light-sheet shows a thinner light distribution over the whole length of the FOV (Fig. 3C). When investigating the sectioned slice of the mouse brain, we can see pronounced image quality improvement in the images with the guided setup (Fig. 3, D and E). The axial resolution decreased towards the right edge of the image leading to reduced contrast and loss of details in the unguided case (Fig. 3F). Meanwhile, the axial resolution was kept similar or even improved towards the right edge preserving contrast and detail over the whole FOV with acoustic LSM (Fig. 3G). The comparison between the images acquired without and with active light manipulation inside the sample (Fig. 3H) highlights the acoustic LSM’s ability to suppress the degradation of axial resolution caused by diffraction of light. Herewith, structures of the mouse’s olfactory bulb like the glomerular layer, mitral cell layer or granule cell layer were observed in better detail.

**Fig. 3.**
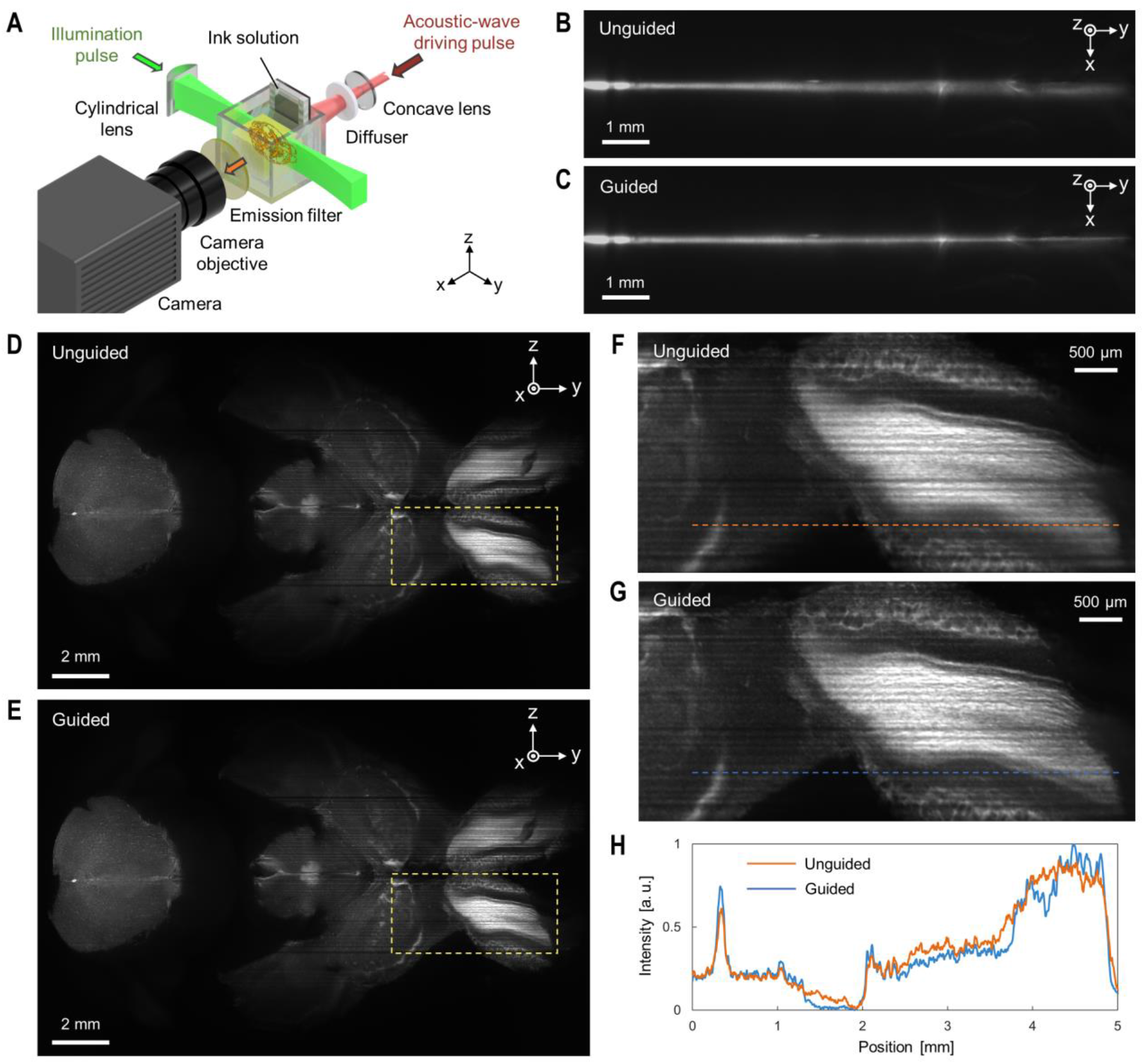
A cleared mouse brain wide-field imaging with acoustic LSFM. **(A)** Schematic diagram of the optical setup for acoustic LSFM. The acoustic wave driving pulse, spatially shaped by a concave lens and diffuser, produces a planar acoustic pulse through absorption at an ink solution. The illumination pulse is cylindrically focused and coupled into the acoustic field inside the sample. Fluorescence images were captured with an emission filter, objective and camera. Light-sheet profiles were captured from the top using an additional optical setup. **(B** and **C)** Light-sheet profile inside the CUBIC cleared mouse brain embedded in hydrogel. The guided light-sheet shows a thinner light distribution. The beam waist of the unguided gaussian beam is located on the left side of the image. **(D** and **E)** Horizontal section of the CUBIC cleared mouse brain captured by a camera objective (×1 magnification, NA = 0.08) with the FOV of 19.3 × 12.4 mm^2^. **(F** and **G)** Enlarged images of boxed regions of (C) and (D), respectively. **(H)** Profiles of fluorescence intensity taken along the orange and blue dotted lines in (F) and (G). The acoustic light-sheet technique shows significant improvements in spatial resolution and image contrast by suppressing the degradation of axial resolution caused by diffraction of light in LSFM.

Increasing the magnification enables a closer look at smaller structures down to single-cell nuclei over the whole FOV. The imaging FOV was 9.7 × 5.9 mm^2^ with a lateral resolution of 2.7 μm, determined by the imaging conditions with the telecentric objective (×2 magnification, NA = 0.12) and the image sensor. In the guided case, the magnified images of the sagittal section of the brain show improved signal-to-noise ratio thanks to the thinner illumination (Fig. 4, A and B). Structures of the anterior commissure fiber tracts (Fig. 4, C and D) and the rostral rhinal vein (Fig. 4, E and F) become clearly visible, which have been masked in the unguided case. When inspecting the mouse brain’s prelimbic area, it is possible to identify small structures and single-cell nuclei when using a guided light-sheet (Fig. 4, G and H). The fluorescence-photon integration in the axial direction of thick light-sheets causes decrease of axial and lateral resolution. This issue limits the identification of detailed structures for a conventional optical imaging method. In our acoustic LSM, the microstructures buried in the background noise in conventional methods can be resolved (Fig. 4, I, J and K) while keeping the wide imaging FOV.

**Fig. 4.**
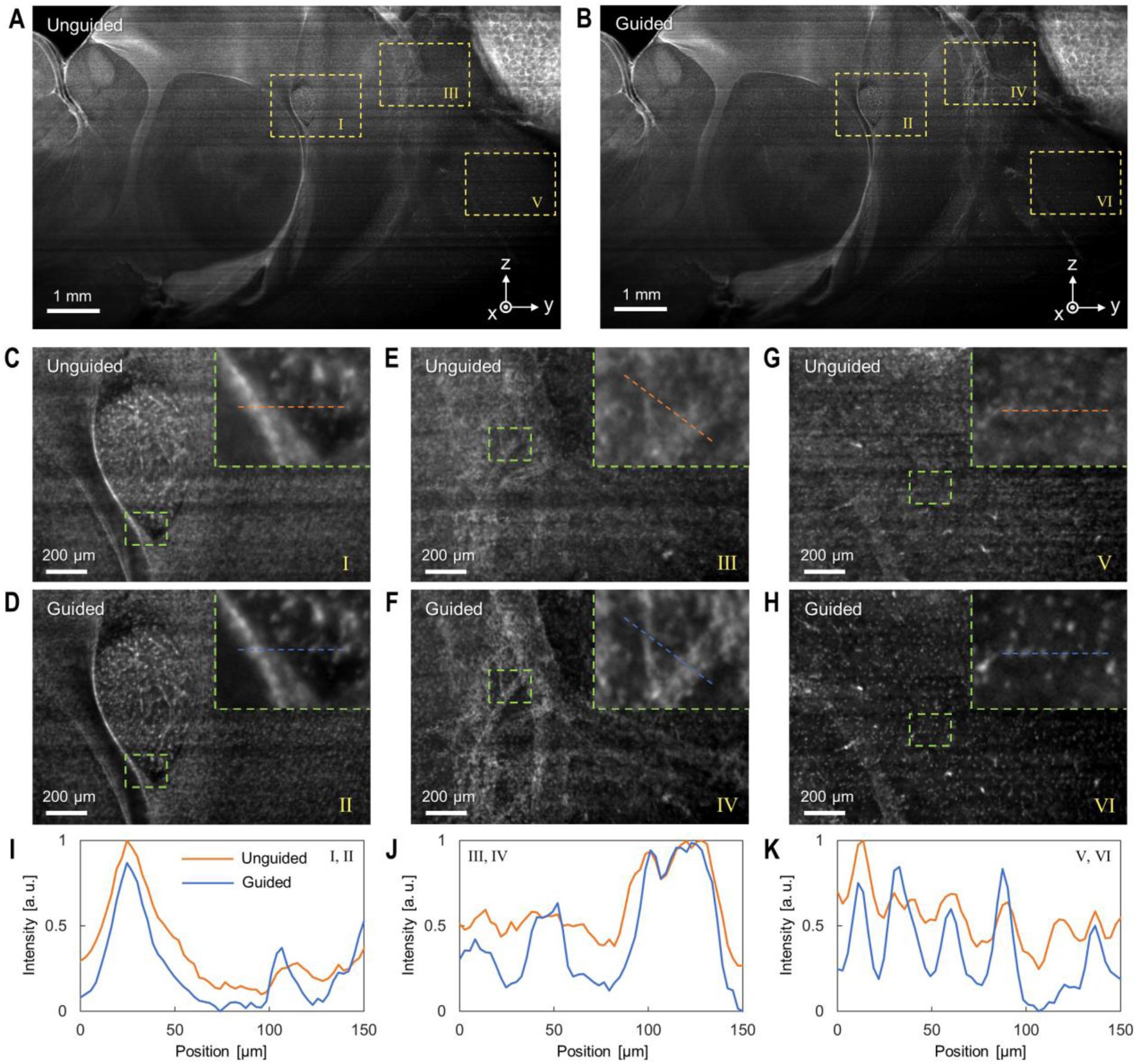
A magnified sagittal section of a transparent mouse brain imaged with acoustic LSFM. **(A** and **B)** Sagittal section of a CUBIC cleared mouse brain, acquired with and without light-guiding. The images were captured by a camera objective (×2 magnification, NA = 0.12) with the FOV of 9.7 × 5.9 mm^2^. **(C** to **H)** Enlarged images of yellow boxed regions I to VI of (A) and (B). Images I, III, V were unguided, while Images II, IV, VI were obtained with acoustic LSFM. The upper right insets in the images are the enlarged images of green boxed regions. **(I**, **J** and **K)** Profiles of fluorescence intensity taken along the orange and blue dotted lines in (C) and (D), (E) and (F), and (G) and (H), respectively. Our technique resolved the detailed structure of the brain tissues (I and J) and imaged the single cells (K) while keeping the wide FOV, both of which were difficult to achieve with the conventional methods.

Acoustic LSM can overcome the fundamental limitation originating from the diffraction of light for conventional LSFM. A single-cell resolved wide-field slice image can be obtained in single-shot which offers reduced photobleaching effects with high image-data throughput for 3D-microscopy applications. Ongoing technology trends for high pixel resolution CMOS sensors and camera array systems (*30*) would boost our approach toward sub-cell-resolution imaging of primate organs. Moreover, we note that the concept of acoustic LSM presented in this report opens the door to a new era in not only wide-field imaging but also the large size 3D printing (*31*), those which require thin light-sheets in their applications.

## Acknowledgments

The authors thank Kenta Kitamura and Chika Shimizu for assisting with experiments.

## Author contributions

K.N. conceived the concept of acoustic LSFM. S.W., A.I. and K.N designed the experiments. S.W. performed the numerical simulation. S.W., A.I., Z.X., H.S., H.Z. and K.N. conducted the basic evaluation of imaging performance. E.A.S. and H.R.U prepared transparent brains, and S.W., A.I. and X.Z. carried out imaging experiments. T.S., I.S., H.F., E.O. and S.T. aided in performing experiments and analyzing the data. H.R.U., S.T. and K.N. supervised the work. S.W. and A.I. wrote the paper and all authors contributed to improvement of the paper.

## Data and materials availability

All data are available in the manuscript or the supplementary materials.

