## Supplementary Text for "Acoustic light-sheet microscopy"

**This PDF file includes:**

Materials and Methods
Supplementary Text
Figs. S1 to S9
Tables S1
Captions for Movies S1 to S2
References (32 - 42)

**Other Supplementary Materials for this manuscript include the following:**

Movies S1 to S2

### Materials and Methods

*Light-sheet confinement by acoustic pulse inside PAA gel sample*

The experimental setup for evaluation of the light-sheet confinement effect inside polyacrylamide (PAA) gel is shown in fig. S6. An acoustic pulse was generated by the photoacoustic driving pulse, generated from a Q-switched Nd:YAG laser source (LS-2132UT, Lotis TII, BY). The center wavelength, pulse duration, and original beam diameter of the pulse are 1064 mm, 5 ns, and 5 mm, respectively. Since the beam of Q-switched laser spreads gradually as it propagates, the photoacoustic driving pulse was focused with a convex lens (f = 500 mm, Thorlabs, US) for shaping the beam diameter to be 5 mm at the position of a photo-absorber. A concave cylindrical lens (f = -40 mm, Thorlabs, US) was placed between the convex lens and the photo-absorber for spatially spreading the beam in the y-direction to produce a wide-planar acoustic pulse. The distance between the convex lens and the concave lens was set to 120 mm and the distance between the concave lens and the photo-absorber was set to 190 mm. The energy of the laser pulse was 101 mJ after the concave lens. The photo-absorber was a carbon ink solution stored inside a container consisting of a glass plate, a silicone frame, and a thin pulse-permeable membrane (Parafilm "M", Bemis, US) attached with an epoxy adhesive. The ink-container and a fluorescent gel sample (see Sample Preparation section for more details) were submerged inside a water-filled quartz glass tank with the size of $100 \times 100 \times 100$ mm$^3$ (SJC-100100, AS ONE, JP). The distance between the sample and the ink container was set to 30 mm.

Another short-pulse Q-switched Nd:YAG laser (GAIA, Rayture Systems, JP) was used to excite fluorescence dyes. The center wavelength and pulse duration of the pulse are 532 nm and 3-5 ns. The beam diameter was expanded to 15 mm and the 3 mm entrance slit was placed before a convex cylindrical lens (f = 100 mm, Thorlabs, US) to create a light-sheet. To observe the confinement of the light-sheet with the acoustic pulse, an sCMOS camera (CS126MU, Thorlabs, USA) equipped with a camera-lens (86574, 50 mm/F1.8, Edmund Optics, US) and a bandpass filter (580BP60, Omega Optical, US) was used. The camera was mounted on a tripod and the fluorescence images were taken from the top. The photoacoustic driving pulse, fluorescence excitation pulse, and the image sensor were synchronized by a digital delay generator (DG535, Stanford Research System, US). Details for the experimental procedure are given in Supplementary note 5.

*Light-sheet confinement by acoustic pulse inside a transparent mouse brain*

The experimental setup for imaging of the transparent mouse brain is schematically shown in fig. S9. For the acoustic pulse generation, the photoacoustic driving pulse (with a center wavelength of 1064 nm, pulse duration of 5 ns, beam diameter of 5 mm and pulse energy of 130 mJ) from a Q-switched Nd:YAG laser source (LS-2132UT, Lotis TII, BY) was spatially spread through a concave lens (f = -75 mm) and homogenized with a diffuser (DGUV10-600, Thorlabs, US) just before irradiating to the carbon ink solution. Similar to the previous section, the ink container consists of a glass plate, on which a silicone frame and a thin pulse-permeable membrane (Parafilm "M", Bemis, US) were attached with an epoxy adhesive. For excitation of fluorescence dyes, a laser pulse (with a center wavelength of 532 nm, pulse duration of 3–5 ns) from a Q-switched Nd:YAG laser source (GAIA, Rayture Systems, JP) was expanded to a diameter of 15 mm and redirected at the sample through a cylindrical lens (f = 100 mm, Thorlabs, US). The whole system was synchronized with a digital delay generator (DG645, Stanford Research Systems, US). Images from the top were captured with an sCMOS image sensor (ORCA-Flash4.0 V3,

Hamamatsu Photonics, JP) with 1× magnification via 2 plano-convex lenses (f = 150 mm, Thorlabs, US) and a 45° mirror. The imaging sections were captured with an additional sCMOS image sensor (ORCA Lightning, Hamamatsu Photonics, JP) equipped with 1× (NA = 0.08) or 2× (NA = 0.12) magnification telecentric objective lens (custom-made product, Shibuya Optical, JP). Both image sensors were equipped with fluorescence filters (#86-111, #84-757, Edmund Optics, US) to cut the light from the fluorescence excitation pulse and the photoacoustic driving pulse.

*Setup for simulation verification with PAA gel sample*

The excitation illumination ($\lambda$ = 532 nm, pulse duration 5 ns, GAIA, Rayture Systems, JP) was created with a horizontally oriented cylindrical lens (f = 200 mm, Thorlabs, US) to generate a thin light-sheet in the x-y plane, that spans over the whole fluorescent gel sample (see Sample preparation section). This kind of illumination can be accurately reproduced in the simulation for comparison. Furthermore, the position of the acoustic pulse is easily detectable this way, and no out-of-focus light masks the light-guiding area. The fluorescence intensity was imaged from the top via a 45° mirror with an sCMOS image sensor (ORCA-Flash4.0 V3, Hamamatsu Photonics, JP) with 1× magnification via 2 plano-convex lenses (f = 150 mm, Thorlabs, US). The acoustic-pulse generation process itself was the same as previously described for the setup for imaging the transparent mouse brain.

### Simulation

The simulation code was implemented in MATLAB (MATLAB R2019b, The MathWorks, Inc., Massachusetts, US), using the Parallel Computing Toolbox. The simulations were executed on an Intel(R) Core(TM) i9-10900X 3.70GHz Processing Unit, with 128 GB RAM and an Nvidia GeForce RTX 2080 SUPER graphic card on the Windows 10 operating system. Details are given in Supplementary note 1, 2, 3, and 4.

### Sample preparation

*Fluorescent PAA gel*

PAA gel containing fluorescent molecules was used to assess the confinement of the light-sheet by the acoustic pulse. PAA gel ($n_0$ = 1.45, $\rho_0$ = 1090 kg/m$^3$, $c_0$ = 1580 m/s) (*32*) was chosen to mimic the CUBIC-cleared tissues samples ($n_0$ = 1.50, $\rho_0$ = 1110 kg/m$^3$, $c_0$ = 1660 m/s). The gel solution was prepared by adding 40% PAA solution and tetramethyl ethylenediamine (TEMED) to water at volume ratio of 25:1:75. The gel solution was stirred and degassed before adding Rhodamine B solution. The final concentration of Rhodamine B was 1 µM. The fluorescent gels solution was poured into a mold (50×50×50 mm$^3$) just before fixation by 0.002% (v/v) of ammonium persulfate. For the simulation verification experiment, the gel size and Rhodamine B concentration were 20 × 20 × 10 mm$^3$ and 10 µM, respectively.

*Transparent mouse brain*

For a demonstration of acoustic LSM on a biological specimen, we used a CUBIC (clear unobstructed brain/body imaging cocktails and computational analysis)-processed mouse brain (*33*). The PFA-fixed mouse brain (C57BL/6N, Japan SLC, Inc., male, 8 weeks old) was delipidated by CUBIC-L (at 37°C for 5 days), stained with propidium iodide [5 µg/mL in 10 mM HEPES buffer (pH 7.5) containing 500 mM NaCl, at 37° for 5 days], and RI-matched with CUBIC-R(N)+

to optically clear the tissue. Before imaging, the cleared tissue was embedded in the CUBIC-R(N)+ reagent with 2% agarose. The sample was immersed in the index-matched CUBIC-R mounting solution (M3294, Tokyo Chemical Industry, JP; $n_0$ = 1.52, $\rho_0$ = 840 kg/m$^3$, $c_0$ = 1272 m/s) during the experiments. This animal experimental procedures and housing conditions of the animals were approved by the Animal Care and Use Committees of the Graduate School of Medicine of the University of Tokyo,

### Image Data Processing

Fluorescence images shown in this paper have been normalized by changing the range of the grayscale values for viewing. In the comparison of the intensity profiles on the line region of interest (ROI) between the unguided images and guided images (Fig. 3H, Fig. 4I, 4J, and 4K), the normalized intensity has been calculated by

$$I_N = \frac{I - \min(I_{Ung\_min}, I_{G\_min})}{\max(I_{Ung\_max}, I_{G\_max}) - \min(I_{Ung\_min}, I_{G\_min})}, \qquad \text{Eq. (1)}$$

where $I_N$ is the normalized intensity of each pixel, $I$ is the original grayscale value in the raw data, $I_{Ung\_max}$, $I_{Ung\_min}$, $I_{G\_max}$ and $I_{G\_min}$ are the maximum values and minimum values on the line ROI in the unguided and guided image data, respectively.

### Supplementary Text

### Supplementary note 1

*Measurements for acoustic pulse properties*

The initial pressure distributions for acoustic simulation in k-Wave were experimentally determined by a polyvinylidene fluoride needle hydrophone (100-100-1, Dr. Muller Instruments, DE) and a digital oscilloscope (DPO 3034, Tektronix, US). For example, an acoustic pulse generated by laser ablation with a laser intensity of 0.06 J/cm$^2$ and a pulse duration of 5 ns in water showed an initial rise distance of 20 µm (fig. S1A). The main positive lobe shows a width of 72 µm, the width is defined here as the distance where the main positive lobe exceeds 10% of the positive pressure peak. The positive and negative pressure-peak values were 428 kPa and -241 kPa, respectively.

The acoustic velocities for the respective media are another important parameter for simulating acoustic pulse propagation and sequence timing in general. Clearing methods like CUBIC, Clarity, or iDisco (*33–35*) use different chemicals for index matching and fixation, of which the acoustic velocities are not known. Therefore, we used the Schlieren imaging method (*36*) for the measurement of acoustic velocities. Acoustic pulses were visualized by a pulsed semiconductor laser source (640 nm Cavilux Smart, Cavitar Ltd, FI) with a pulse duration of 20 ns, two lenses (f = 50 and mm f = 100 mm, Thorlabs, US), and a high-speed camera (HPV-X2, Shimazu, JP) equipped with a camera objective (Laowa 60mm f/2.8 2X Macro, Venus Optics, CHN), to capture the fast dynamics of the acoustic pulse with 5 million frames per second (Movie S1 and fig. S2).

By measuring the pixel-displacement per time interval in the acquired high-speed video, the acoustic velocity of the CUBIC-cleared mouse brain and CUBIC-R matching oil were 1669 $\pm$ 10 m/s ($N$ = 5) and 1272 $\pm$ 14 m/s ($N$ = 11), respectively.

### Supplementary note 2

*Acoustic pulse propagation simulation*

Since biological samples are usually not in liquid form, it is difficult to measure the pressure profile inside the sample. Moreover, the acoustic profile inside water, which can be measured by the hydrophone, differs from that inside the sample. This originates from sound speed differences among the media. Therefore, the acoustic pulse propagation was simulated to estimate the pressure profile inside the sample.

The acoustic pulse propagation was simulated with the open-source, third-party-MATLAB toolbox k-Wave (*37*). The toolbox is based on the k-space pseudospectral method and is designed for time-domain acoustic and ultrasound simulations in tissue-like media. It describes fluid dynamics with a series of coupled first-order partial differential equations, including momentum conservation, mass conservation, and pressure-density relation. First, we initialized the computational grid and defined the acoustic properties for the media (sample medium, ink-container, and others). The laser-ablation event is realized in the form of a time-varying pressure source. The ablation area is defined via a source mask and weighted with an exponential decay function to approximate the medium's optical absorption. Furthermore, the ablation mask was weighted with a gaussian profile similar to the spatial profile of the ablation laser. The temporal pressure source followed a smoothed Friedlander curve, as measured with the hydrophone. To keep the computation time low, the computational grid was initialized asymmetrically. Due to acoustic pulse's planar nature, the pressure profile does not rapidly change perpendicular to the propagation direction. Hence, the grid step size perpendicular to the propagation direction can be chosen larger while keeping the simulation accurate. After investigating several media with different acoustic parameters, we concluded that light-guiding efficiency does not vary drastically (fig. S4). Therefore, acoustic LSM should be equally suited for soft-tissue-like media with different acoustic properties. The exact simulation conditions can be seen in Table S1.

### Supplementary note 3

*Pressure-Refractive Index relation*

After obtaining the waveform at the position of interest in the acoustic simulation, the pressure profile was converted to the refractive index. In general, the refractive index $n$ is a function of the density $\rho$ of the medium. With the help of the Lorentz-Lorenz equation, the density can be related to the local refractive index (*38*). The equation is given by

$$n = \sqrt{\frac{2Q\rho + 1}{1 - Q\rho}}, \qquad \text{Eq. (2)}$$

where $Q$ the molar refractivity, which can be determined by inserting the reference refractive index $n_0$ and the density $\rho_0$ at reference pressure. Furthermore, the relationship of pressure and density of a fluid in adiabatic compression processes can be described by an equation of state with the form

$$P_a = \frac{\rho_0 c^2}{\gamma}\left[\left(\frac{\rho}{\rho_0}\right)^{\gamma} - 1\right], \qquad \text{Eq. (3)}$$

where $P_a$ stands for the pressure of the medium, $\rho$ for density, $c$ is its acoustic velocity, and $\gamma$ is a material parameter, which is usually chosen as $\gamma = 7$ for nearly incompressible fluids such as water and hydrogels (*39*). In literature, this or similar forms of the equation of state are often referred to as Cole equation of state or Tait equation of state (*40*). The estimated refractive-index contrasts for a selected number of pressure amplitudes are shown in fig. S1B. Notable is the asymmetric shape, with a steeper pressure gradient at the wavefront.

### Supplementary note 4

*Optical simulation with FFT-BPM*

The spatial dimensions of the positive peak of laser-induced waveguides are generally in the micrometer domain. Therefore, light-guiding properties become hard to interpret by simple raytracing due to the properties of gaussian beams. The 2D Fast Fourier transform Beam Propagation Method (FFT-BPM) provides a fast and efficient way to simulate electromagnetic wave propagation in waveguide-like structures and is well-suited for planar acoustic waveguides (*26*). It is derived from the scalar wave equation under the slowly varying envelope approximation, assuming propagation is mostly parallel along the waveguide axis, with the refractive index varying around a reference index $n_0$. BPM is based on Maxwell's equation, and the fundamental problem is written and solved in the form of a scalar wave-equation (Helmholtz-equation), as described in (*27*). The equation is stated as

$$\frac{\partial^2 u}{\partial y^2} + 2i\beta\frac{\partial u}{\partial y} + \frac{\partial^2 u}{\partial x^2} + \frac{\partial^2 u}{\partial z^2} + (k^2 - \beta^2)u = 0, \qquad \text{Eq. (4)}$$

where $k^2 = k_0^2 n^2$ , $\beta^2 = k_0^2 n_0^2$ and $k$ is the wavenumber, with the $k_0 = 2\pi/\lambda_0$ in free space. $n$ is the refractive index of the waveguide and $n_0$ is the reference refractive index of the medium. Applying the small-angle approximation, assuming wave-propagation dominantly in the y-direction, and only considering the 2D-case, the problem can be simplified to

$$\frac{\partial u}{\partial y} = \frac{i}{2\beta}\left(\frac{\partial^2 u}{\partial x^2} + (k^2 - \beta^2)u\right). \qquad \text{Eq. (5)}$$

This parabolic partial differential equation was numerically solved using a 2-step FFT algorithm (*26, 27*), where the input field is first propagated a small distance $\Delta y$ using the FFT, and then inverse Fourier-transformed and phase-corrected according to the refractive index distribution. These steps are repeated to obtain the field at any given finite distance.

FFT-BPM only simulates one-way instead of rigorously solving the Maxwell-equations in all directions at all positions of the computational grid at the same time. This makes FFT-BPM less computationally expensive than the acoustic pulse-propagation of k-Wave. Additionally, due to the slowly varying envelope approximation assumption, a much coarser grid step size in the y-direction can be chosen, which reduces the computational load further. This enabled the interpolation of the refractive index profile obtained in the acoustic propagation simulation to a finer grid in the x-direction for more accurate optical simulations while keeping computation time small.

Experimental and simulation results show good agreement in shape and light intensity with each other (fig. S3). The background was subtracted for both images to better visualize the relative change of fluorescence intensity due to optical guiding. The full width at half maximum (FWHM) of the light-guiding area was $53.6 \pm 2.6$ µm ($N$ = 5) for experimental results and 38.1 µm for simulation (fig. S3B).

### Supplementary note 5

*Measurements of the length of the acoustic pulse*

The length of the generated acoustic pulse used in the light-sheet confinement evaluation with PAA was measured using the Schlieren method. For length estimation, the acoustic pulse was measured in pure water. The optical components for acoustic pulse generation were the same as those described in the Materials and Methods section, except for that the concave cylindrical lens (f = -40 mm, Thorlabs, US) was turned by 90 degree (fig. S7A). The illumination pulse was generated by a pulsed semiconductor laser source (640 nm Cavilux Smart, Cavitar Ltd, FI) to visualize the acoustic pulse in water. The illumination pulse duration was selected as 10 ns. One lens (f = 100 mm, Ø75 mm) was utilized to collimate the light and another lens (f = 200 mm, Ø75 mm) was used to focus the light. The sCMOS camera (CS126MU, Thorlabs, US) equipped with a camera-lens (#86574, 50 mm/F1.8, Edmund Optics, US) was used for imaging. A bandpass filter (#65229, CWL = 640 nm, FWHM = 10 nm, GO Techspec, Edmund Optics, US) and a ND filter (NE02A-A, Thorlabs, US) were set in front of the camera lens. A digital delay generator (DG535, Stanford Research System, US) was utilized to synchronize the whole system.

Schlieren images were taken when the acoustic pulse was generated. Additionally, a background image was also taken when there was no acoustic pulse. To exclude the background noise, the background image data was subtracted from the original image. From the Schlieren imaging result of the acoustic pulse, it can be observed that a planar acoustic pulse was generated by the photoacoustic driving pulse (fig. S7B). To estimate the length of the planar acoustic pulse, the full width at half maximum (FWHM) of the acoustic pulse wavefront was measured. This FWHM should not be confused with the FWHM of the thickness of the wavefront, previously mentioned (Supplementary notes 4). The values are first smoothed by 5 point moving average and then normalized to the maximum value. The dotted line in fig. S7B shows the half maximum. While there are a few points whose intensity are lower than 0.5 within the middle region, the FWHM is estimated as 33.7 mm. This value is consistent with the results in the PAA light-sheet confinement experiment, where the confinement effect was confirmed up to 30 mm.

### Supplementary note 6

*Light-sheet confinement by an acoustic pulse inside PAA gel sample*

Fluorescence images were captured at different delay times between the photoacoustic driving pulse and the illumination pulse (fig. S8A). Delay time *t* is the delay between photoacoustic driving pulse and illumination pulse. The negative *t*-values correspond to time delays, where the acoustic wavefront did not yet arrive at the light-sheet plane when the illumination pulse was generated. The positive *t*-values correspond to time delays, where the acoustic wavefront had already passed the exact position of the light-sheet plane when the illumination pulse was generated. At $t = 0$ ns, the light-sheet was optimally coupled with the acoustic pulse. The confinement of the light-sheet was observed when the acoustic pulse had reached the position of the light-sheet (Movie S2). The full-width half-maximum (FWHM) of the light-sheet (imaged from top) was calculated along the light propagation direction. The guided light-sheet's FWHM showed a reduction of up to 80.7% at y = 29.3 mm compared to a non-guided light-sheet. Fig. S8B shows the change of the intensity profiles with time, along x-axis at y =29.3 mm. The intensity was normalized to the maximum. It can be observed that the light-sheet was confined at $t = 0$ ns.

### Supplementary note 7

*3D Image acquisition*

In conventional LSFM imaging for large samples, high-resolution 3D image data is acquired by positioning the beam waist of the light-sheet at a certain position and acquire a stack of images by scanning the sample perpendicular to the light-sheet in x-direction, generating a subvolume. After acquiring one stack, the sample is translated in y-direction to the next position to acquire the next subvolume. This is repeated until the whole sample is scanned and the subvolumes are registered and combined. With *m* steps in x-direction and *n* light-sheet repositioning in y-direction, the total image acquisition time of the 3D volume can be written as **m × n × exposure time + m × n × stage step time**.

New developments for 3D LSFM imaging try to minimize the amount of stage moving time, as it is usually the bottleneck for fast 3D image acquisition (*41*,*42*). One way to reduce imaging time was presented in the form of a tiling light-sheet, where the position of the light-sheet is adjusted with phase modulation (*41*). That way, for each scanning step in x-direction, the whole image plane can be acquired without moving the sample. Without the sample translations in y-direction, the image acquisition time reduces to **m × n × exposure time + m × stage step time.**

Another method to reduce the stage moving time was presented in form of a scan termed MOVIE (*42*), in which the stage moves continuously in a uniform motion, removing the acceleration and deceleration time of the stage. The image acquisition procedure itself is similar as in conventional LSFM imaging. The total imaging acquisition time is given by
**n × x-scanning time.**

In acoustic LSFM, no repositioning of the light-sheet is necessary, as the whole image-plane is captured in a single-shot fashion. As the acoustic pulse generation is usually similar or faster than the stage step time, the total image acquisition time is given by **m × exposure time + m × stage step time**. Theoretically, if sufficient timing and synchronization can be achieved, acoustic LSFM should be compatible with the MOVIE sequence. This would further reduce the image acquisition time to just **x-scanning time.**

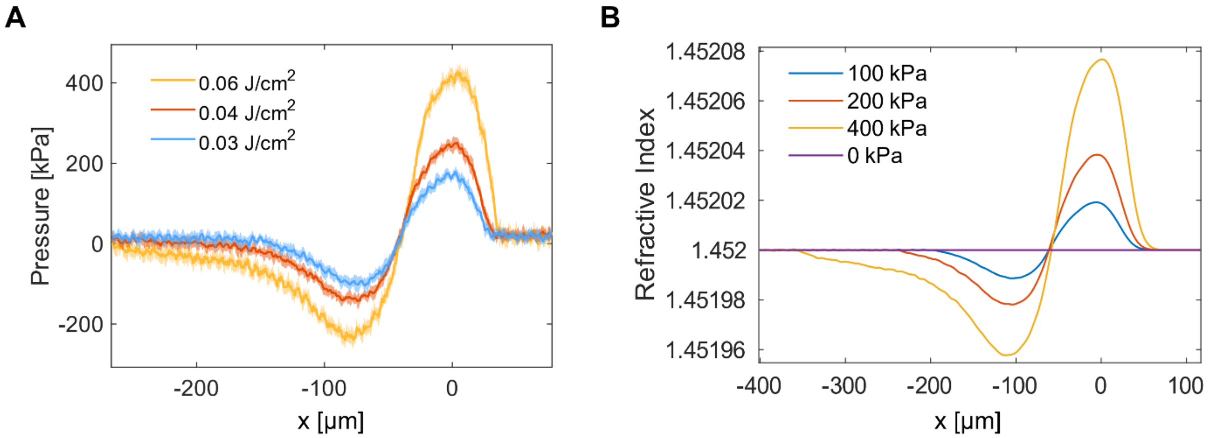


**Fig. S1 Acoustic profiles of laser-induced acoustic pulse and estimated refractive index. (A)** Acoustic profiles of laser-induced acoustic pulses inside water measured by a hydrophone over a range of laser intensities. The hydrophone was placed 10 mm away from the ablation point. **(B)** Estimated refractive indices from (A). The legend indicates the positive pressure amplitude used for the initial pressure input in the simulation.

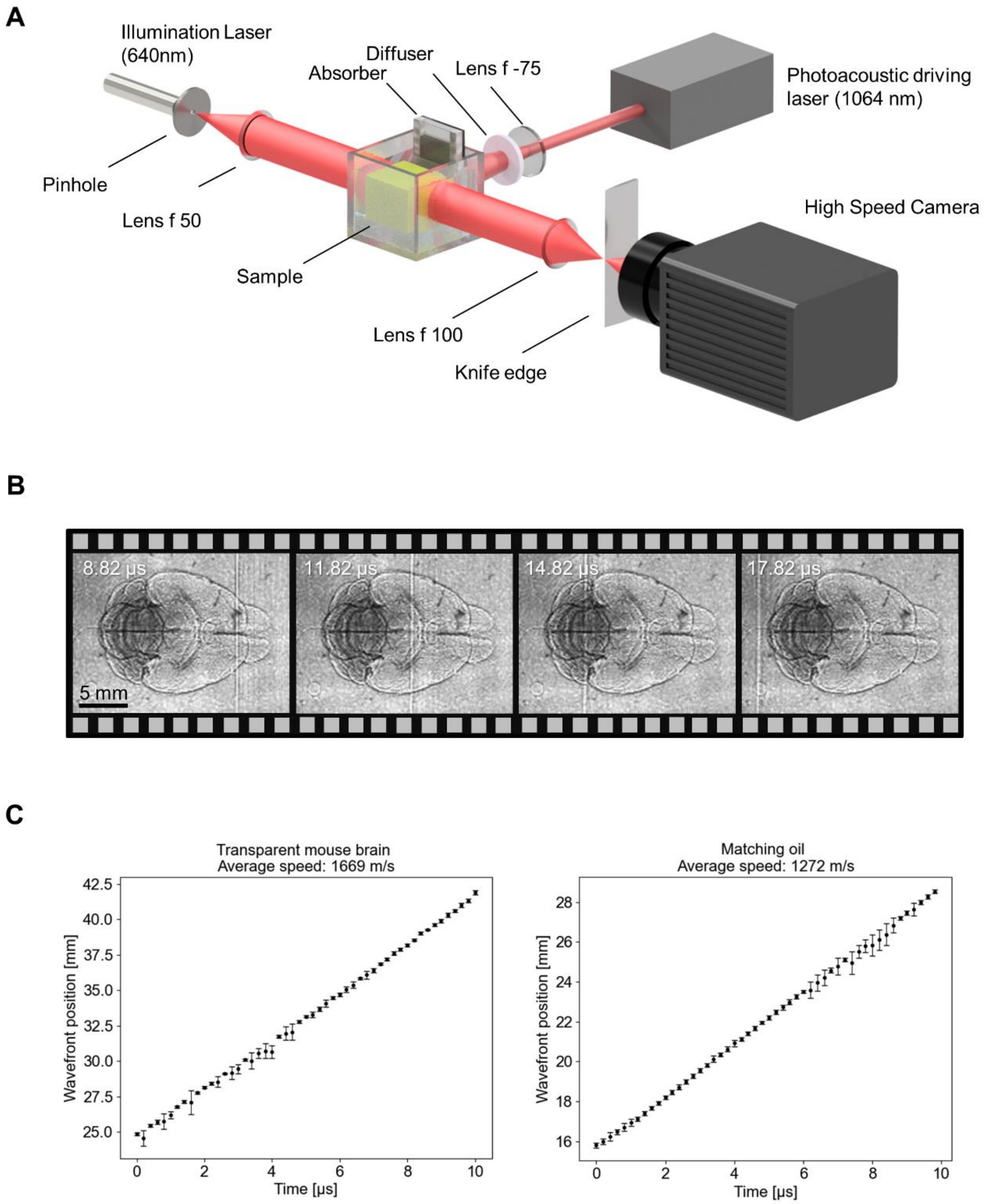


**Fig. S2 Optical Schlieren measurement of the acoustic pulse propagating inside a CUBIC-cleared mouse brain sample. (A)** Schematic diagram of the experimental setup for Schlieren imaging. (**B)** Representative images of acoustic pulse propagation in the sample, which were used to estimate the acoustic velocity. (**C)** By measuring the pixel-displacement per time interval in the acquired high-speed video, the acoustic velocities of 1669 $\pm$ 10 m/s ($N$ = 5) for the transparent mouse brain and 1272 $\pm$ 10 m/s ($N$ = 11) for the matching oil were estimated. The time interval between the video frames was 200 ns.

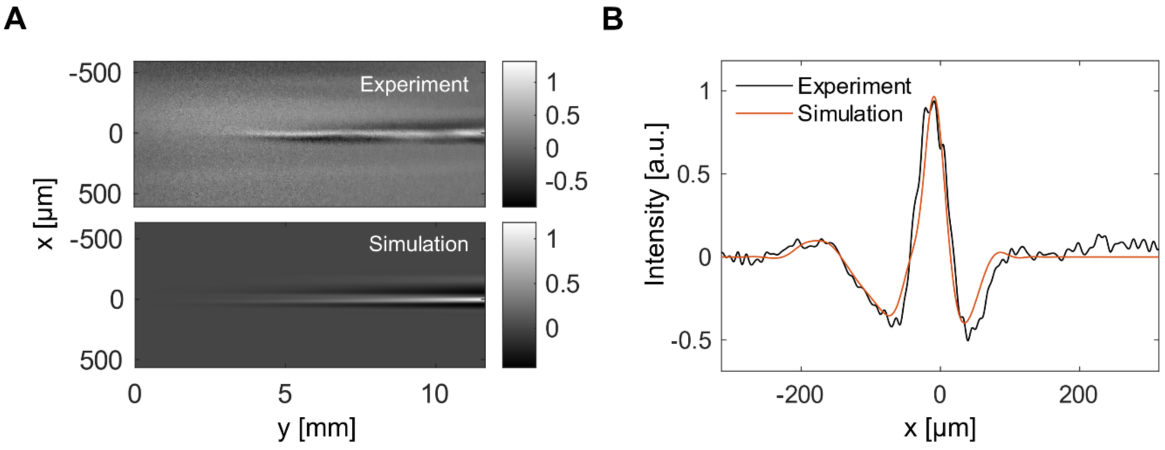


**Fig. S3 Comparison of experimental and simulation results for light-guiding with an acoustic pulse. (A)** Experimental and simulation results for the visualized light-guiding area by focusing a horizontal light-sheet through the orthogonally oriented acoustic pulse in a PAA gel sample containing Rhodamine B. **(B)** Corresponding fluorescence intensity profiles at y = 11 mm for experimental and simulation results.

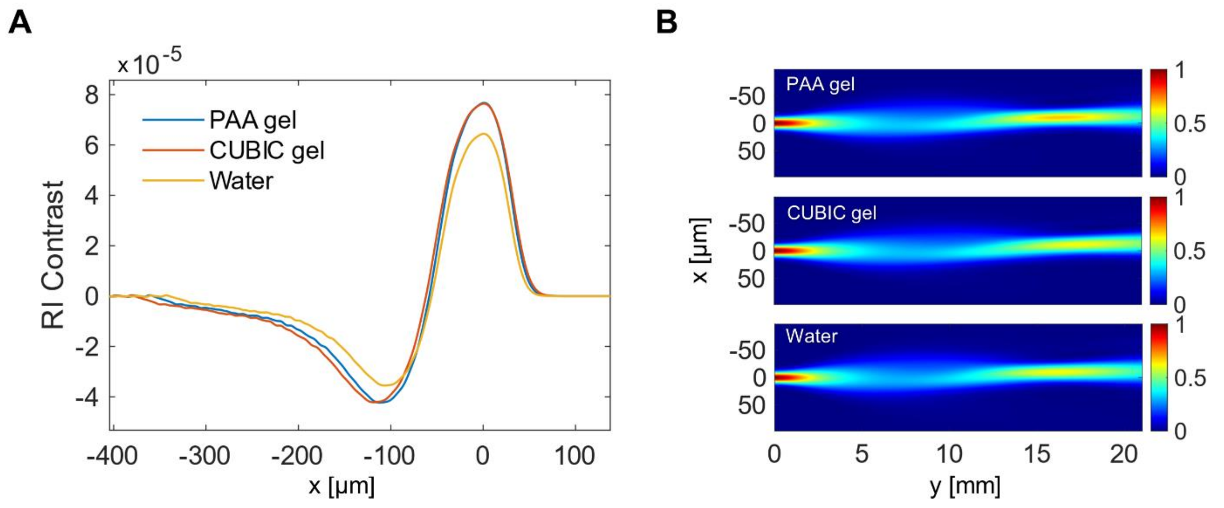


**Fig. S4 Simulated light-guiding with different media. (A)** Results for the simulated refractive index contrasts produced by laser-induced acoustic pulses for different media. The media investigated are PAA gel ($n_0$ = 1.45, $\rho_0$ = 1090 kg/m$^3$, $c_0$ = 1580 m/s), a CUBIC gel sample ($n_0$ = 1.5, $\rho_0$ = 1110 kg/m$^3$, $c_0$ = 1669 m/s) and water ($n_0$ = 1.33, $\rho_0$ = 1000 kg/m$^3$, $c_0$ = 1500 m/s). The refractive index contrasts produced by laser-induced acoustic pulse did not drastically differ for different media. (**B)** Light-guiding simulation results for different media. Due to the relatively similar refractive index contrast, the light-guiding efficiency only differed slightly.

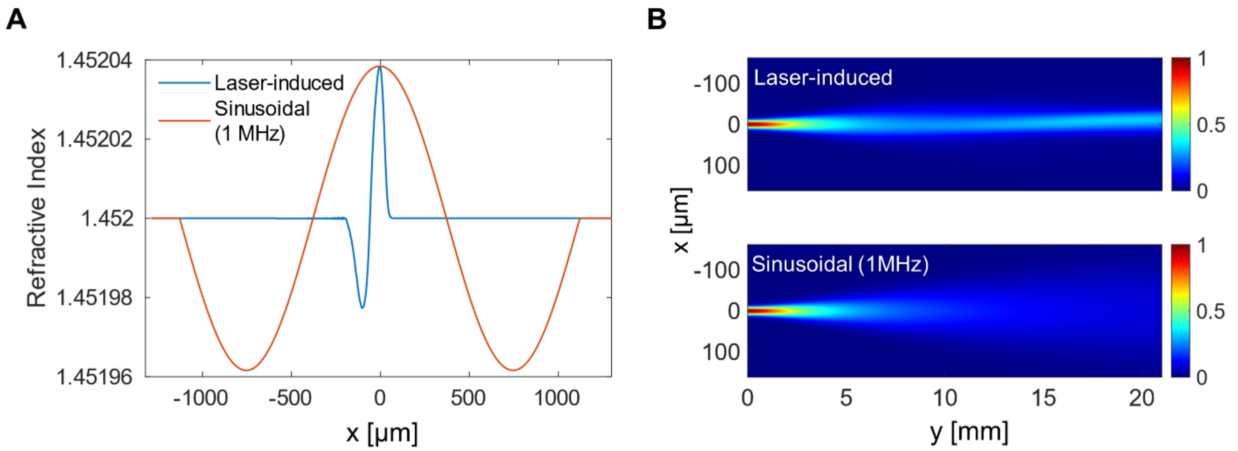


**Fig. S5 Simulated light-guiding for the laser-induced acoustic pulse and a 1 MHz sinusoidal pulse. (A)** Simulated refractive indices for the laser-induced acoustic pulse and 1 MHz sinusoidal pulse. The positive peak-pressure was set to 200 kPa. **(B)** Simulated light-guiding for the laser-induced acoustic pulse and the 1 MHz pulse. The light guiding by laser-induced acoustic pulse showed higher light-guiding power than the 1 MHz pulse with the same pressure amplitude.

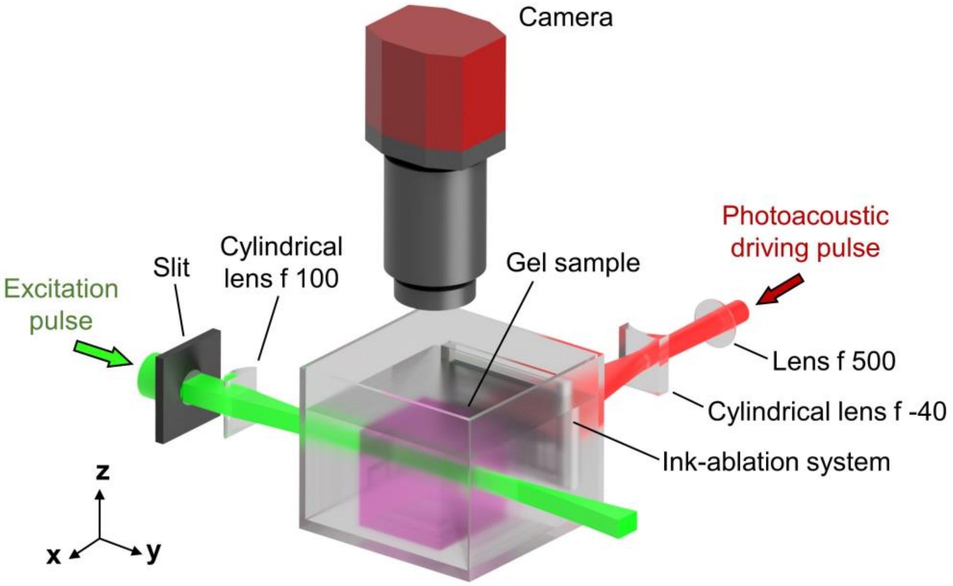


**Fig. S6 Schematic of the experimental setup for evaluating the light-sheet confinement effect inside the PAA gel sample.** A photoacoustic driving pulse with a center wavelength of 1064 nm was shaped and directed to a carbon ink photo-absorber. A fluorescent PAA gel sample was irradiated with an excitation light-sheet with a center wavelength of 532 nm. The PAA gel sample and the photo-absorber were immersed in a water tank. For visualizing the light-sheet confinement effect, fluorescence images were captured by an sCMOS camera from the top.

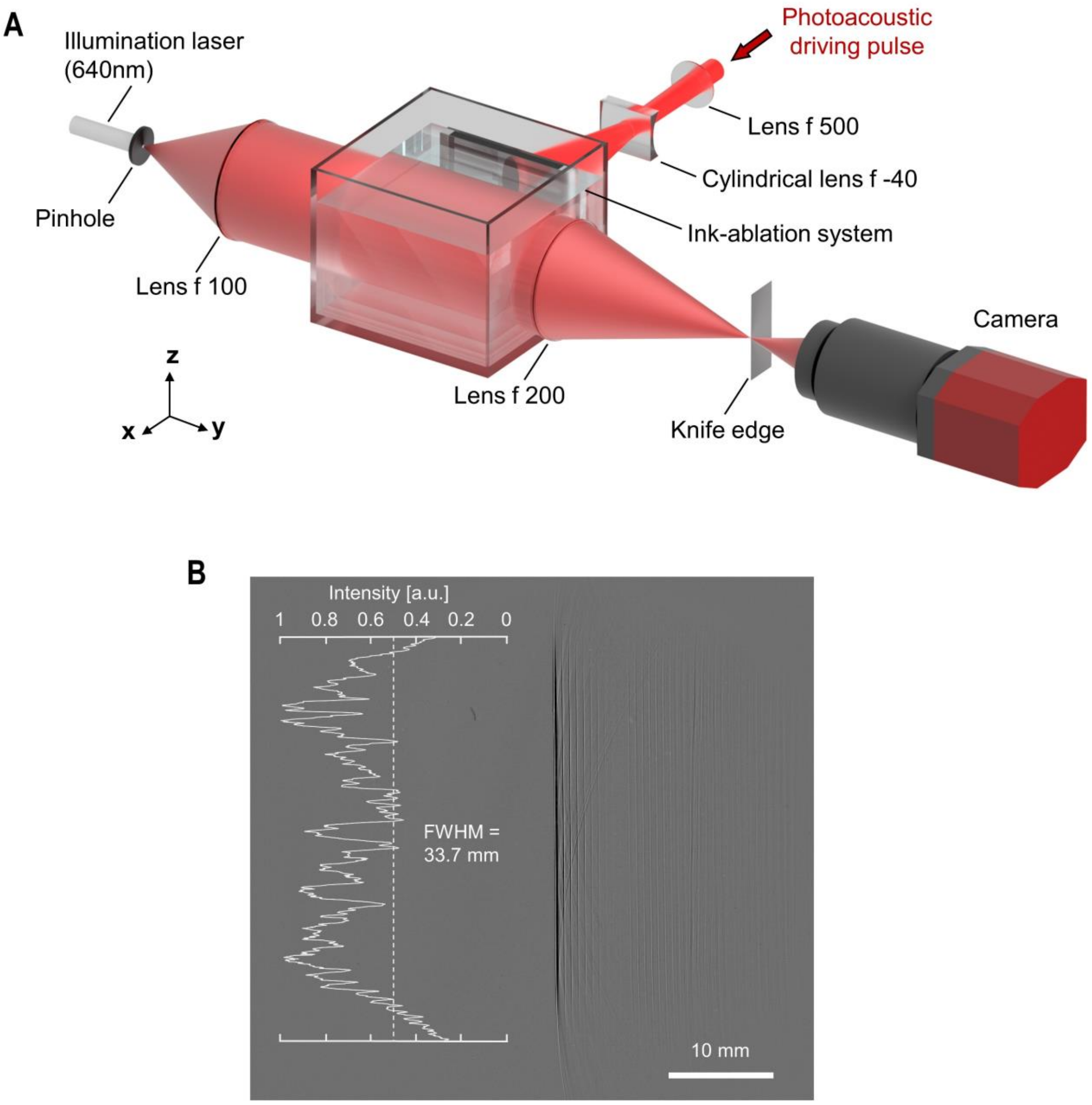


**Fig. S7 Measurement of the length of the acoustic pulse used in light-sheet confinement evaluation with PAA by Schlieren imaging. (A)** Schematic diagram of the experimental setup for Schlieren imaging of large scale acoustic pulses. **(B)** Representative Schlieren image of acoustic pulse propagation in water after background subtraction. The maximum intensities of the acoustic pulse wavefront are shown on the left. Values are smoothed by 5 point moving average and then normalized to the maximum. The dotted line shows the half maximum.

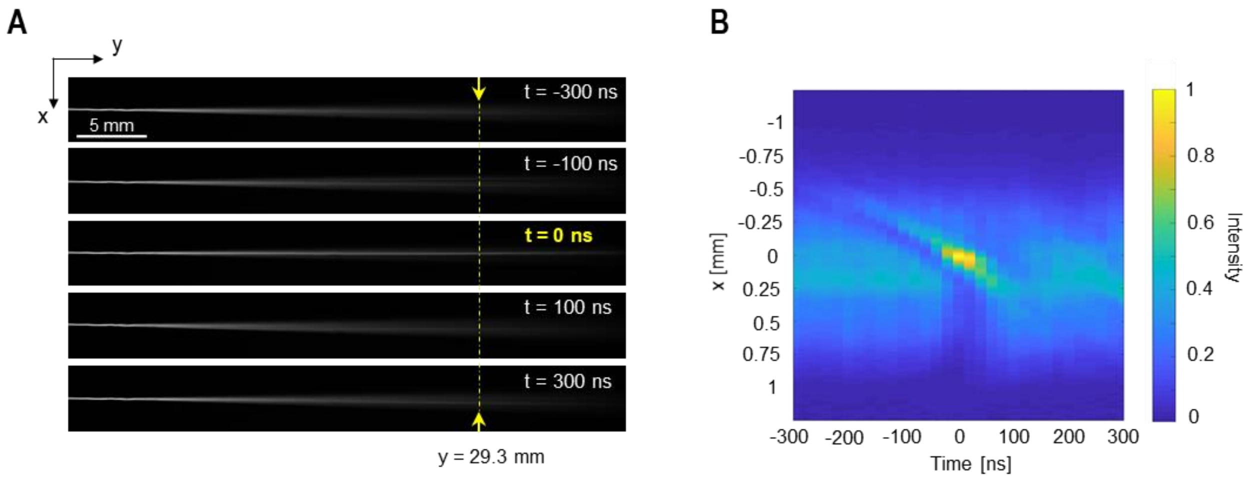


**Fig. S8 Dynamics of light-sheet confinement by an acoustic pulse inside a PAA gel sample containing Rhodamine B. (A)** Representative fluorescence images of the light-sheet confinement by an acoustic pulse at different time delays. The illumination light-sheet pulses were focused at 2.5 mm from the left edge of the images. At $t = \pm 100$ ns and $\pm 300$ ns, the light-sheet was diverging along the $y$-direction because the acoustic pulse was not accurately coupled to the light-sheet. At $t = 0$ ns, the light-sheet was confined along the $y$-direction. **(B)** Transient fluorescence intensity at $y = 29.3$ mm. At this position, the FWHM was reduced by a maximum of 80.7% with guiding, compared to a non-guided light-sheet.

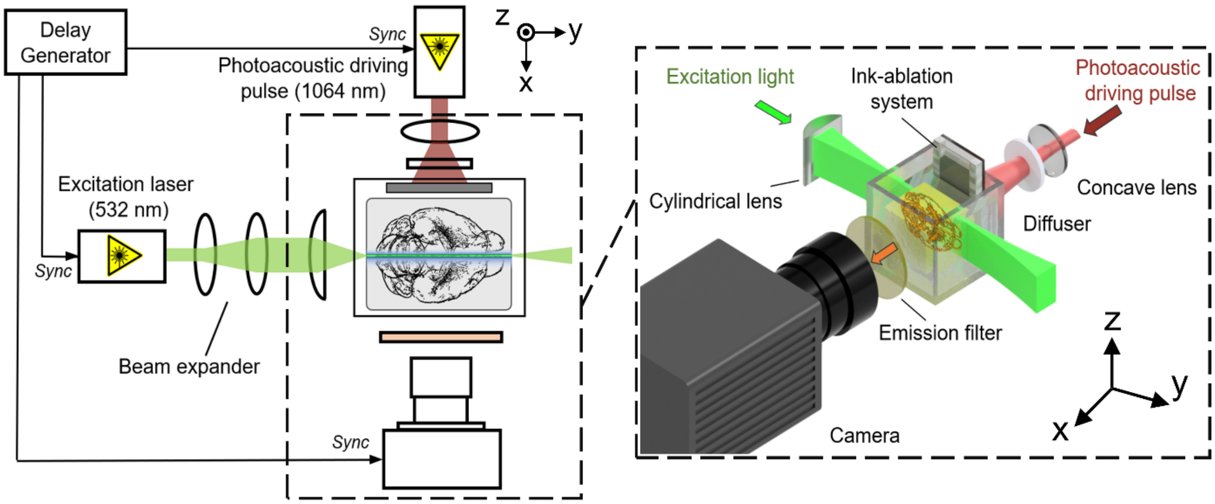


**Fig. S9 Experimental setup for transparent brain imaging.** Schematic diagram of the optical setup for acoustic LSFM. Light-sheet profiles were captured from the top using a 45° mirror, emission filters, and an additional image sensor which are not shown in this illustration.

**Table S1** Parameters for acoustic simulation in k-Wave

| | Water | PAA gel | CUBIC gel |
|---|---|---|---|
| Acoustic velocity $c_0$ [m/s] | 1500 | 1580 | 1669 |
| Density $\rho_0$ [kg/m$^3$] | 1000 | 1110 | 1110 |
| Number of grid points (x × y) | | 2000 × 1000 | |
| Grid spacing in x [μm] | | 2.5 | |
| Grid spacing in y [μm] | | 20 | |
| Temporal resolution [ps] | | 250 | |

**Movie S1 Propagating acoustic pulse in CUBIC-cleared mouse brain.** The bright line in the center of the image is the planar acoustic pulse propagating through the sample. The wavefront was followed by periodic low-pressure acoustic pulses which are reflected acoustic pulses from the glass plate of the absorber.

**Movie S2 Motion pictures of light-sheet confinement by acoustic pulse inside the PAA gel sample.** The upper-right part is the schematic diagram of the acoustic pulse propagating towards the light-sheet. The timing $t = 0$ ns denotes the time stamp, when the light-sheet was confined by the acoustic pulse to reduce the FWHM 80.7% at y = 29.3 mm.
